# Plasmo3Net: A Convolutional Neural Network-Based Algorithm for Detecting Malaria Parasites in Thin Blood Smear Images

**DOI:** 10.1101/2024.12.12.628235

**Authors:** Afolabi J. Owoloye, Funmilayo C. Ligali, Ojochenemi A. Enejoh, Oluwafemi Agosile, Adesola Z. Musa, Oluwagbemiga Aina, Emmanuel T. Idowu, Kolapo M. Oyebola

**Affiliations:** Center for Genomic Research in Biomedicine (CeGRIB), College of Basic and Applied Sciences, Mountain Top University, Kilometer 12, Lagos-Ibadan Expressway, Nigeria; Parasitology and Bioinformatics Unit, Department of Zoology, Faculty of Science, University of Lagos, Lagos, Nigeria; Nigerian Institute of Medical Research, Lagos, Nigeria; Biochemistry Department, Faculty of Basic Medical Science, University of Lagos, Lagos, Nigeria; Genetics, Genomics and Bioinformatics Department, National Biotechnology Research and Development Agency, Abuja, Nigeria

**Keywords:** Malaria, Deep Learning, Convolutional Neural Network, Plasmo3Net, Diagnostic Workflow

## Abstract

Early diagnosis of malaria is crucial for effective control and elimination efforts. Microscopy is a reliable field-adaptable malaria diagnostic method. However, microscopy results are only as good as the quality of slides and images obtained from thick and thin smears. In this study, we developed deep learning algorithms to identify malaria-infected red blood cells (RBCs) in thin blood smears. Three algorithms were developed based on a convolutional neural network (CNN). The CNN was trained on 15,060 images and evaluated using 4,000 images. After a series of fine-tuning and hyperparameter optimization experiments, we selected the top-performing algorithm, which was named Plasmo3Net. The Plasmo3Net architecture was made up of 13 layers: three convolutional, three max-pooling, one flatten, four dropouts, and two fully connected layers, to obtain an accuracy of 99.3%, precision of 99.1%, recall of 99.6%, and F1 score of 99.3%. The maximum training accuracy of 99.5% and validation accuracy of 97.7% were obtained during the learning phase. Four pre-trained deep learning models (InceptionV3, VGG16, ResNet50, and ALexNet) were selected and trained alongside our model as baseline techniques for comparison due to their performance in malaria parasite identification. The topmost transfer learning model was the ResNet50 with 97.9% accuracy, 97.6% precision, 98.3 % recall, and 97.9% F1 score. The accuracy of the Plasmo3Net in malaria parasite identification highlights its potential for automated malaria diagnosis in the future. With additional validation using more extensive and diverse datasets, Plasmo3Net could evolve into a diagnostic workflow suitable for field applications.

## 1.0 INTRODUCTION

Malaria is a deadly disease caused by parasites transmitted to humans through the bite of infected female *Anopheles* mosquitoes. Early diagnosis of the disease is crucial to decreasing the number of malaria victims. Microscopy is considered the gold standard for malaria diagnosis and quantification [1]. However, malaria diagnosis can be enhanced by utilizing digital image processing techniques to reliably detect parasites in images of red blood cells (RBCs) from microscope slides [2, 3]. The advent of computer vision and convolutional neural networks (CNNs) has enabled successful image classification tasks [4-6].

The connectivity patterns between neurons in a CNN mirrors the organization of the visual cortex, where individual cortical neurons respond to stimuli only within a restricted receptive field of the visual field [7]. This biological principle inspired the hierarchical structure of learning layers in a CNN model [8]. CNNs effectively utilize the spatial correlations of visual patterns, such as edges in images, to reduce the number of training parameters [8, 9]. This improves the accuracy of the feedforward-backpropagation training procedure. As such, CNNs provide a general-purpose learning framework that eliminates the need for substantial feature extraction and fine-tuning. This advantage over traditional classification methods allows deep learning models to effectively capture highly complex data patterns [10].

The early applications of Convolutional Neural Networks (CNNs) date back to the 1990s for tasks such as speech recognition [11] and text recognition [12], and later expanded to handwriting recognition [13] and natural image recognition [14]. With the advent of ImageNet [15], CNN’s image classification performance received significant recognition. AlexNet also achieved a top-five error rate of 15.3% on the ILSVRC-2012 competition, which had 10,184 categories and 8.9 million images [15]. Subsequent improvements steadily reduced the top-five error rate, from 14.8% by ZFNet in 2013 [16] to 6.7% by GoogLeNet in 2014 [17] and finally to 3.6% by ResNet50 in 2015 [5]. These advancements have demonstrated the powerful capabilities of CNNs for large-scale image recognition tasks.

Previous approaches for computer vision-based classification of *Plasmodium*-infected and uninfected RBCs have mostly focused on archived image sets [18]. While the outcomes reported in these studies have been promising, it is crucial to demonstrate the robustness and performance of these methods on field datasets. The existing deep-learning models also have considerable room for improvement in this area. In this study, we explored a customized deep-learning algorithm using a convolutional neural network (CNN) architecture and evaluated its robustness and performance metrics on a dataset of blood smear images of *P. falciparum* isolates from Nigeria.

## 2.0 METHODOLOGY

The dataset used in this study included images of Giemsa-stained slides of thin blood smears obtained from 48 *Plasmodium falciparum*-infected and 23 healthy individuals during a previous antimalarial therapeutic efficacy study [19]. Images were digitized using a light microscope x100 resolution. An iPhone 14 with a high-resolution digital camera (4032 x 3024 pixels) was used to capture each field. The experimental workflow included dataset acquisition, image preprocessing, image segmentation, data augmentation, and building a deep learning model based on convolutional neural network (CNN) architecture (Figure 1). To determine the performance of the model on unseen data, we downloaded *P. falciparum* microscope slide images from the National Institutes of Health (NIH) [18, 20]. The dataset contained two classes - parasitized RBC images and uninfected RBC image cell images. Each class had 13,794 images. When analyzing the dataset, we performed manual curation to enhance accurate sample labelling. We used 12,200 manually curated images of the parasitized and uninfected classes. Moreover, to check the performance of our model, several transfer learning models such as AlexNet, ResNet50, Inception V3, and VGG16 were trained and evaluated for comparison.

**Figure 1:**
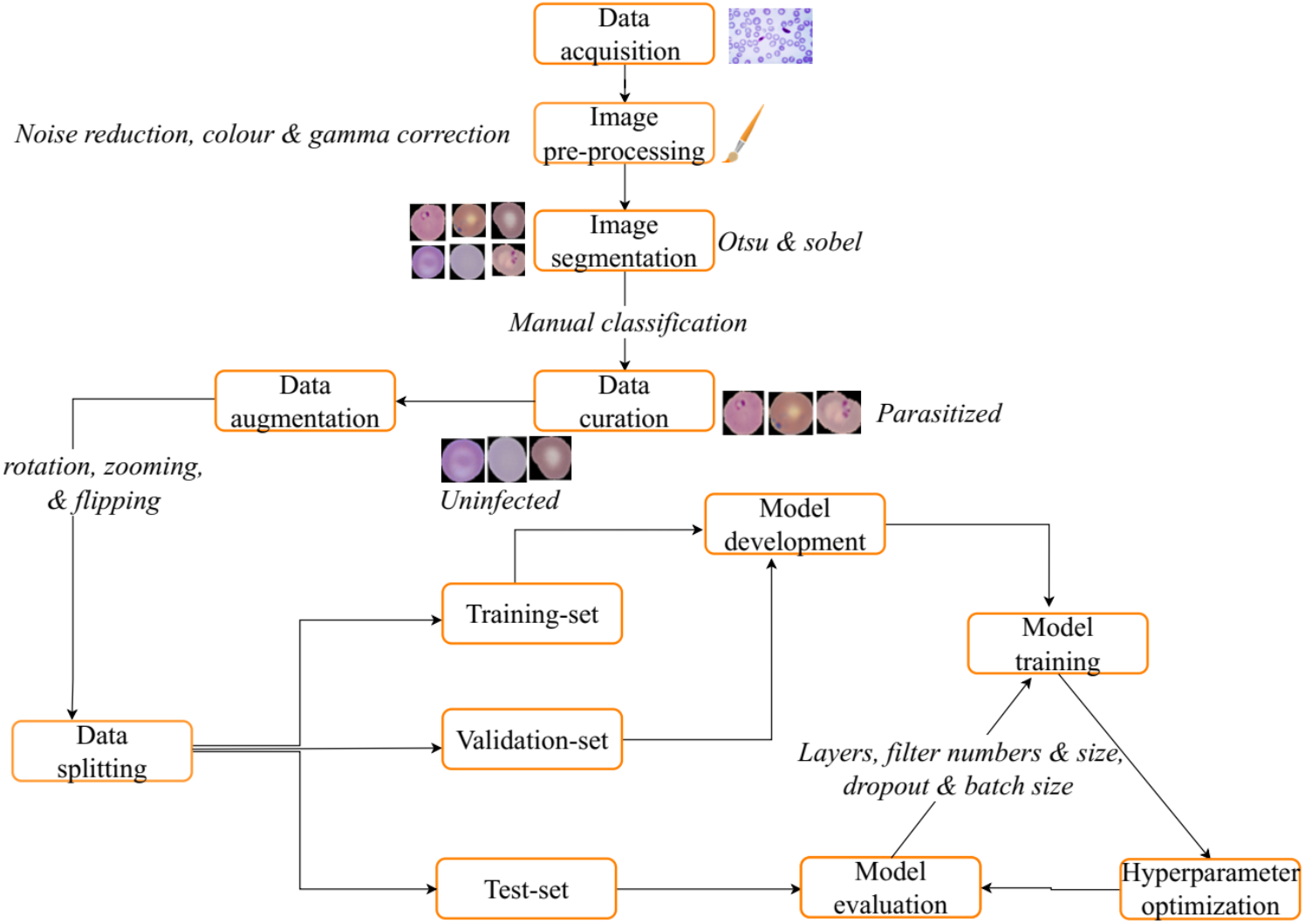
Experimental workflow for developing the convolutional neural network

### 2.1 Image preprocessing

#### 2.1.1 Cropping

We cropped specific regions of interest within images (we excluded the dark area of the slide image). The coordinates of the crop are contained in four specified values: X, Y, width, and height. The X or the horizontal coordinate of the crop region was the left side of the image where the crop started. The vertical coordinate of the crop region, the y-coordinate, was defined as the upper edge from which cropping begins. The specified width and height parameters specified the rectangular crop area, or the extent of the complete area for extraction 2450 x 2450 pixels.

#### 2.1.2 Noise reduction

A bilateral filter was used to remove noise from digital images while preserving edges (Figure 2). The bilateral filter is a non-linear technique to preserve edges by applying less smoothing to pixels that are different in color from the central pixel while applying more smoothing to similar pixels [21]. We applied the bilateral filter to the input image using the “cv2.bilateralFilter()” function provided by the OpenCV library in Python. Subsequently, a filtering neighborhood diameter of nine pixels, a sigma value of 75 in the color space, and a sigma value of 75 in the coordinate space were configured. These parameter values were determined to effectively balance noise reduction and edge preservation for the image dataset.

**Figure 2.**
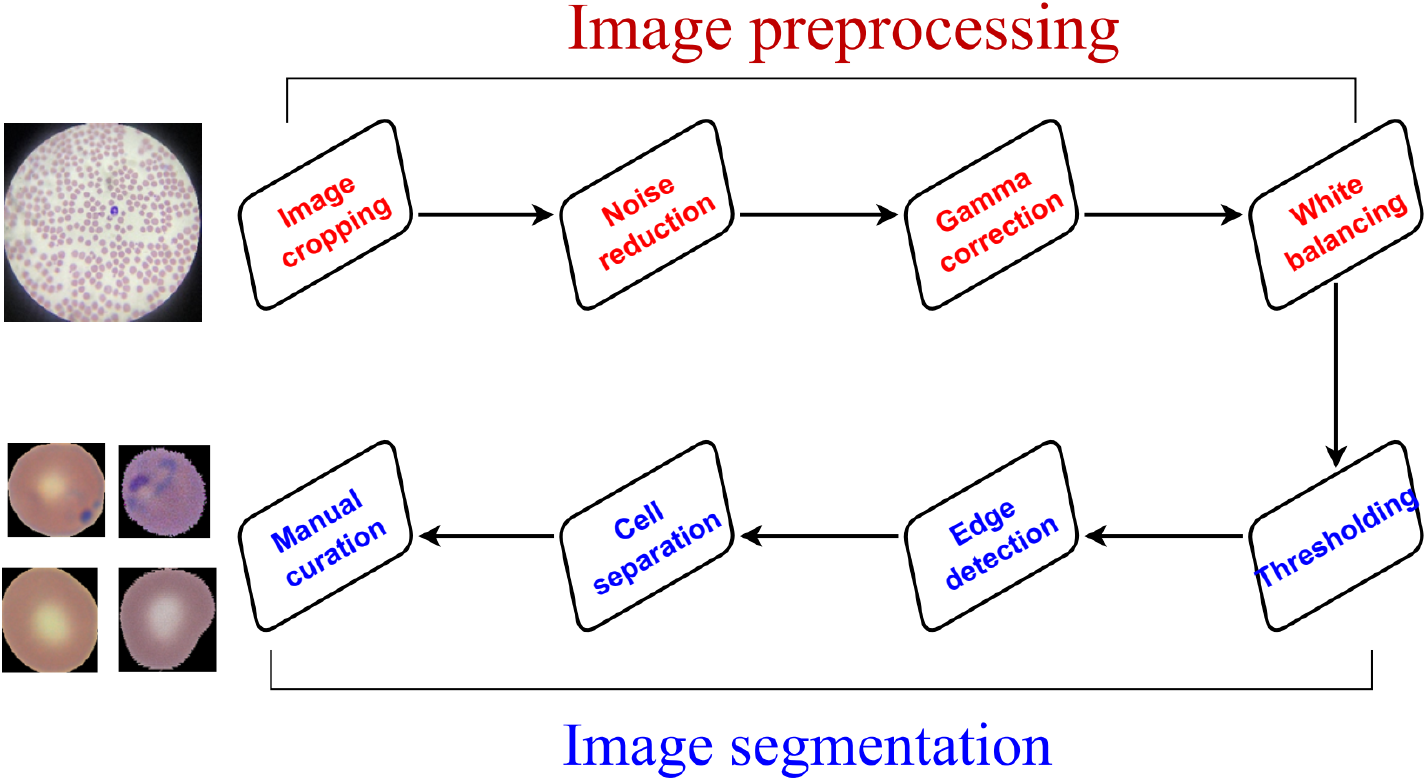
Image preprocessing and segmentation workflow.

#### 2.1.3 Gamma correction

Gamma correction was applied to enhance the brightness and contrast of the digital images. Gamma correction selectively modified the luminance information of the images while preserving the original hue and saturation values. The images were then converted from the default BGR color space to the hue, saturation, value (HSV) color space. Subsequently, we determined the desired “mid-tone” brightness of 0.8 and calculated the gamma value based on the channel’s mean pixel value (V). The gamma value was applied to the V channel of the HSV images. The V channel was then merged with the HSV channels and converted back to the BGR color space. The flexibility to adjust the gamma value based on the image characteristics allows for further optimization and fine-tuning of the desired visual outcomes.

#### 2.1.4 Color correction (white balancing)

Uninfected RBCs occasionally contain artefacts or remains of other types that a deep learning classifier could incorrectly identify as *Plasmodium* parasites [22]. Thus, color normalization was required before classification. The white balancing operation reduced the image’s dynamic range while enhancing the overall contrast and brightness to achieve a better visual quality of content. It was used to improve the visual quality and analyze digital images in different image processing applications (such as computer vision and remote sensing).

White balance was achieved by multiplying the input image by a scaling factor of 1.2 full-size image. The scaling factor of 1.2 used in this analysis was an optimal value obtained because of the characteristics of the input image and the desired visual outcome. This was intended to increase the overall brightness of the image. The scaled image was divided by the maximum value along the height and width dimensions of the original image.

### 2.2 Image Segmentation

The segmentation stage involved the separation of the foreground (RBCs, WBCs, and other blood components) against the background by determining the threshold value that divided the image into two groups based on intensity [18, 23]. The images were then converted from the BGR colour space to grayscale to simplify the segmentation process. We then transformed the image into a binary image preprocessing [24]. The Otsu approach automatically selected a threshold level that optimized the variation between the two histogram classes of the grayscale images. This step resulted in a binary image that highlighted the segmented RBCs. The segmented cells’ contours were detected using the “*findContours()*” function. The Sobel edge detection algorithm was applied to compute the gradients of the input images in the horizontal and vertical directions [25]. Subsequently, the gradients were combined to form a single-edge map highlighting the images’ edges and boundaries. To enhance the effectiveness of segmentation techniques and leverage the potential of color picture segmentation, we adopted the procedure described by Diaz *et al*. [22] concentrated on identifying malaria by applying the moving algorithm k-means clustering in conjunction with the hue, saturation, and intensity (HSI) colour space to obtain full segmentation of RBC (both infected and uninfected).

### 2.3 Data augmentation and splitting

Data augmentation was applied to the training dataset to enhance accuracy [26]. This generated 14,980 images (7,490 each for infected and uninfected RBC) after which 4080 images were selected and augmented by random rotation, random zooming, vertical flipping, and horizontal flipping, making a total of 19060 images. The test dataset comprised 4,000 randomly selected images (2,000 each for infected and uninfected), while the remaining 15,060 images (7,530 each for infected and uninfected) were split (80:20) into training and validation datasets.

### 2.4 Plasmo3Net Architecture

A deep learning algorithm was developed using CNN architecture with 13 layers: three convolutional, three max-pooling, one flatten, four dropouts (5% each), and two fully connected layers (Figure 3). Rectified Linear Unit (ReLU) activation function was used for this configuration. An input image size of 64 × 64 pixels with three channels was sufficient to capture the neighborhood context for a final decision. The initial conv2D layer had 32 filters and a kernel size of 3 × 3. Meanwhile, the subsequent convolutional layer filters were 64, 128, and 128. Following each Conv2D layer, a MaxPool2D layer with a pool size of (2, 2) was applied to down-sample the feature maps. A dropout layer of 5% was then added after the MaxPool2D layer to prevent overfitting. The dropout layer discarded 5% of the neurons. A flatten layer was introduced to convert the 2D output from the previous convolutional layers into a 1D tensor. Following the flatten layer, a fully connected layer with 128 units and ReLU activation was incorporated. A dropout layer with a rate of 5% was included to prevent further overfitting. The final layer of the model produced the classification output, a fully connected layer with 1 unit and a sigmoid activation function. This layer provided a probability for binary classification. The binary cross-entropy loss function was used to calculate the error between predicted actual and predicted output. The model was then compiled using the Adam optimizer [27], an efficient variant of stochastic gradient descent. Furthermore, the model performance was evaluated using the performance metrics. Moreover, two other architectures were developed; a 16-layer model consisting of four convolutional layers, four max-pooling layers, one flatten, five dropouts, and two fully connected layers, and a 19-layer architecture consisting of five convolutional, five max-pooling, one flatten, six dropouts, and two fully connected layers. The two models were also trained, finetuned, and optimized alongside the Plasmo3Net. However, the best CNN architecture (PlasmoNet) was chosen after experimenting with and fine-tuning the hyperparameters. Models were trained using a Dell Alienware Aurora R13 with Nvidia GeForce 3090, 12th Gen intel Core^TM^ i9, 64 GB DDR5 ram, and 24 threads running on Ubuntu 20.04.6 LTS 64-bit OS.

**Figure 3:**
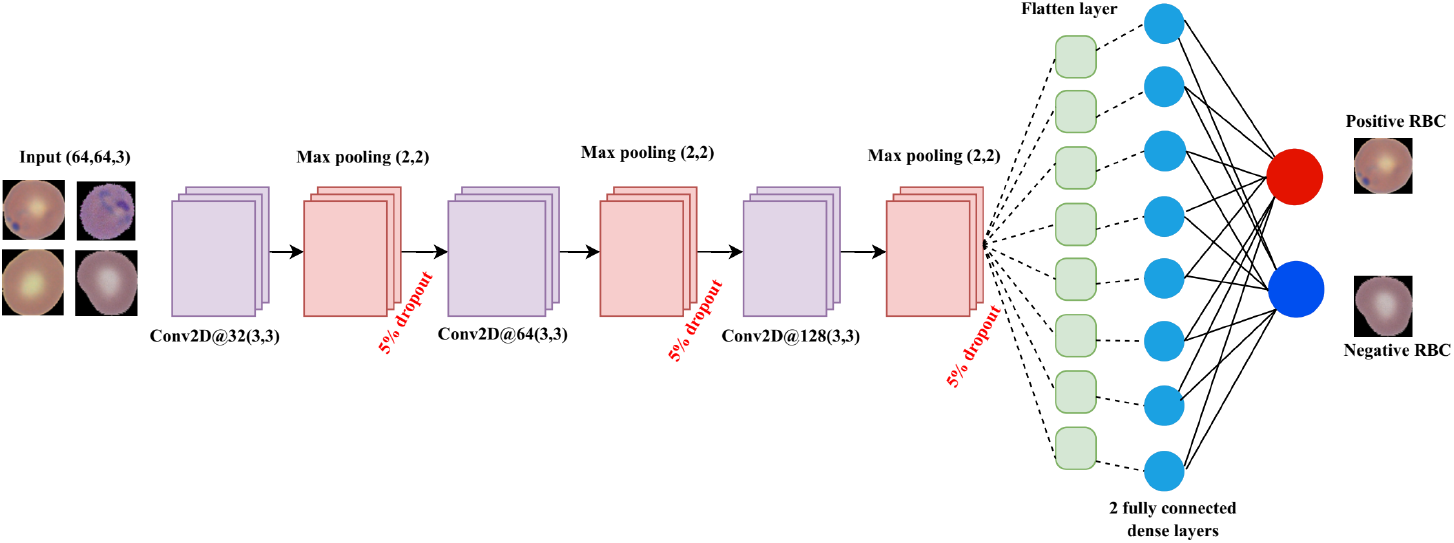
Plasmo3Net convolutional neural network architecture

### 2.5 Transfer learning

Four pre-trained deep learning models - InceptionV3, VGG16, ResNet50, and AlexNet – were used to validate Plasmo3Net. The pre-trained and proposed Plasmo3Net models had the same dataset of 15,060 images (7,530 parasitized and 7,530 uninfected cell images). For binary classification, all models’ final layers were modified to a dense layer with a sigmoid activation function rather than a softmax. The Visual Geometry Group (VGG16) architecture [28] contained 16 layers of convolution network and three fully connected layers with 138 million trainable parameters. The network received an image with a fixed size of (125x125); hence, the matrix’s shape was 125,125,3. The kernel had a stride of one pixel and a size of 3×3. To improve the model’s classification, a max pooling was applied over a 2x2 pixel window with a stride size of two, and then a rectified linear unit (ReLU) was used. The final layer has a softmax activation function changed to sigmoid as the model was meant for binary classification. The AlexNet pre-trained architecture comprises five convolutional layers and three fully connected layers [15]. The first convolutional layer has 96 kernels of size (11, 11) with a stride of 4, followed by an overlapping max-pooling layer with a kernel size of 3×3 and stride two. The overlapping occurred because the stride size was lesser than the kernel size. Max-pooling was adopted to preserve the depth of the image array while reducing its height and width. There were two more max-pooling layers and four convolutional layers. In addition, there were two fully connected layers with 4096 units each. The Softmax function was changed to sigmoid as the model was meant for a binary classification task. The ReLU function was applied to speed up the model after every layer (fully connected and convolutional). The Residual Network (ResNet50) architecture had 50 layers (48 convolutional layers, 1 average-pooling layer, and 1 max-pooling layer) with 23.5 million trainable parameters and 53 thousand non-trainable parameters [5]. ResNet50 addressed deep neural network vanishing and gradient problems using shortcuts or skip connections to create residual blocks. It comprised five steps, each with a convolutional and identity block (each convolutional block has three convolutional layers, and each identity block has three convolutional layers). The core building block of ResNet50 was the residual block, which allowed the network to learn the residual mapping between the input and output of the block. Convolutional layers, inception modules, auxiliary classifiers, and fully connected layers with 23.8 million trainable parameters constituted the Inception v3 architecture [29]. The inception v3 network began with a convolutional layer followed by a max-pooling layer that hosted several inception modules, at the heart of the architecture. Each inception module consisted of many parallel convolutional layers with different filter sizes. Concatenating the outputs of these parallel convolutions allowed features of various scales to be learned simultaneously by the network. The Inception modules also utilized 1x1 convolutions to reduce the input channel number, thus decreasing computational complexity. ReLU was the primary activation function used by convolutional layers, inception modules, and fully connected layers. A softmax activation function was employed in the final layer of the network to output class probabilities for classification tasks. Nevertheless, it was replaced with sigmoid when binary classification was carried out. We trained all the models with a batch size of 64 for 25 epochs. The model performance during the training and validation process was evaluated using the accuracy and loss.

## 3.0 RESULTS

The accuracy and loss curve for training and validation was plotted for each model to understand the rate of improvement in the model performance epoch (Figure 4). The Plasmo3Net had the least trainable parameters with zero non-trainable parameters (Table 1), making it train faster than the other two CNN models and the four pre-trained architectures. The Inception V3 had the highest number of trainable parameters (23.8 million). ResNet50, AlexNet, and VGG16 had total trainable parameters of 23.5 million, 21.5 million, and 15.2 million, respectively. In contrast, the Plasmo3Net has a total number of 683,329 trainable parameters.

**Table 1:** Convolutional Neural Network Model Parameters.

| Model | Trainable parameter | Non-trainable parameter |
| --- | --- | --- |
| AlexNet | 21,585,281 | 0 |
| Inception V3 | 23,868,578 | 34,432 |
| ResNet50 | 23,534,592 | 53,120 |
| VGG16 | 15,239,489 | 0 |
| 16-layer CNN | 913,729 | 0 |
| 19-layer CNN | 1,831,745 | 0 |
| Plasmo3Net | 683,329 | 0 |

**Figure 4.**
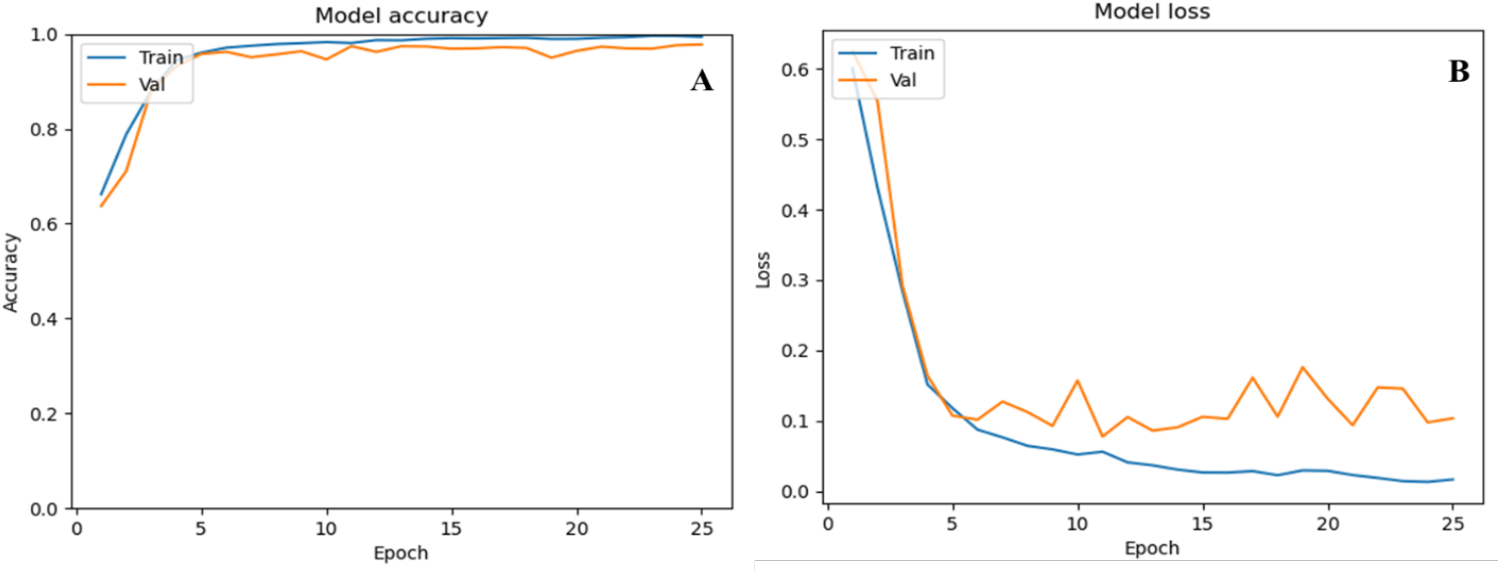
Plasmo3Net learning curve: (A) accuracy, (B) loss.

Comparing the Plasmo3Net with other transfer-learning models, the Plasmo3Net had the best training and validation accuracy (Figures 4 and 5). The discrepancy between the validation and training accuracy indicated that our model had not been over-fitted. By experimenting with various batch sizes and number of epochs, we discovered a trade-off between training speed and the convergence behavior of the model. We achieved a stable gradient estimate through optimization batch size (64). For the least cost function to converge smoothly, this stability was necessary. Consistent updates to the model parameters were made possible by the less noisy gradients produced by the adjusted batch size.

**Figure 5.**
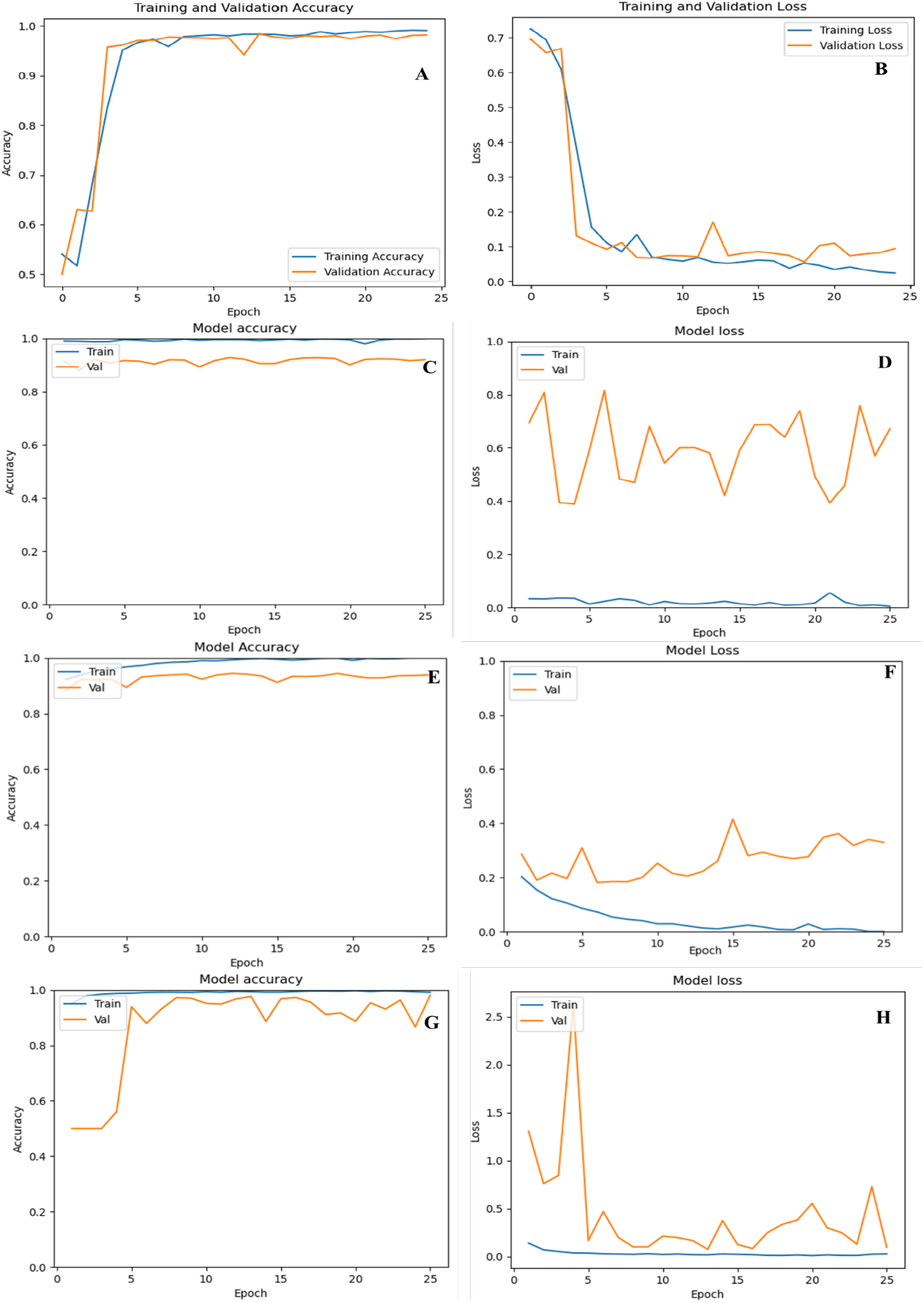
Pre-trained model learning curve: (A) VGG-16 accuracy, (B) VGG-16 loss, (C) Inception V3 accuracy, (D) Inception V3 loss, (E) ResNet50 accuracy (F) ResNet50 loss (G) AlexNet accuracy (H) AlexNet loss.

The maximum training accuracy of 99.5% and validation accuracy of 97.7% were obtained during the learning phase of the Plasmo3Net. Less variation exists between training and validation accuracy, indicating that the model could not have been over-fitted. The AlexNet architecture reached a maximum training accuracy of 99.8% and validation accuracy of 91.9%, however, the validation loss had an average of 65%. The training and validation loss of the Plasmo3Net decreased steadily from the first epoch to the last epoch during the training process, while in the transfer learning models, the validation either fluctuated or increased.

The Plasmo3Net and the transfer learning models were evaluated during the training phase by making predictions and fitting the model to the unexposed data from the test dataset. Finally, the optimal model was implemented on the test dataset (Figure 6). Their performance metrics examined the model potential with parameters like accuracy, F1 score, recall, precision, sensitivity, and specificity.

**Figure 6.**
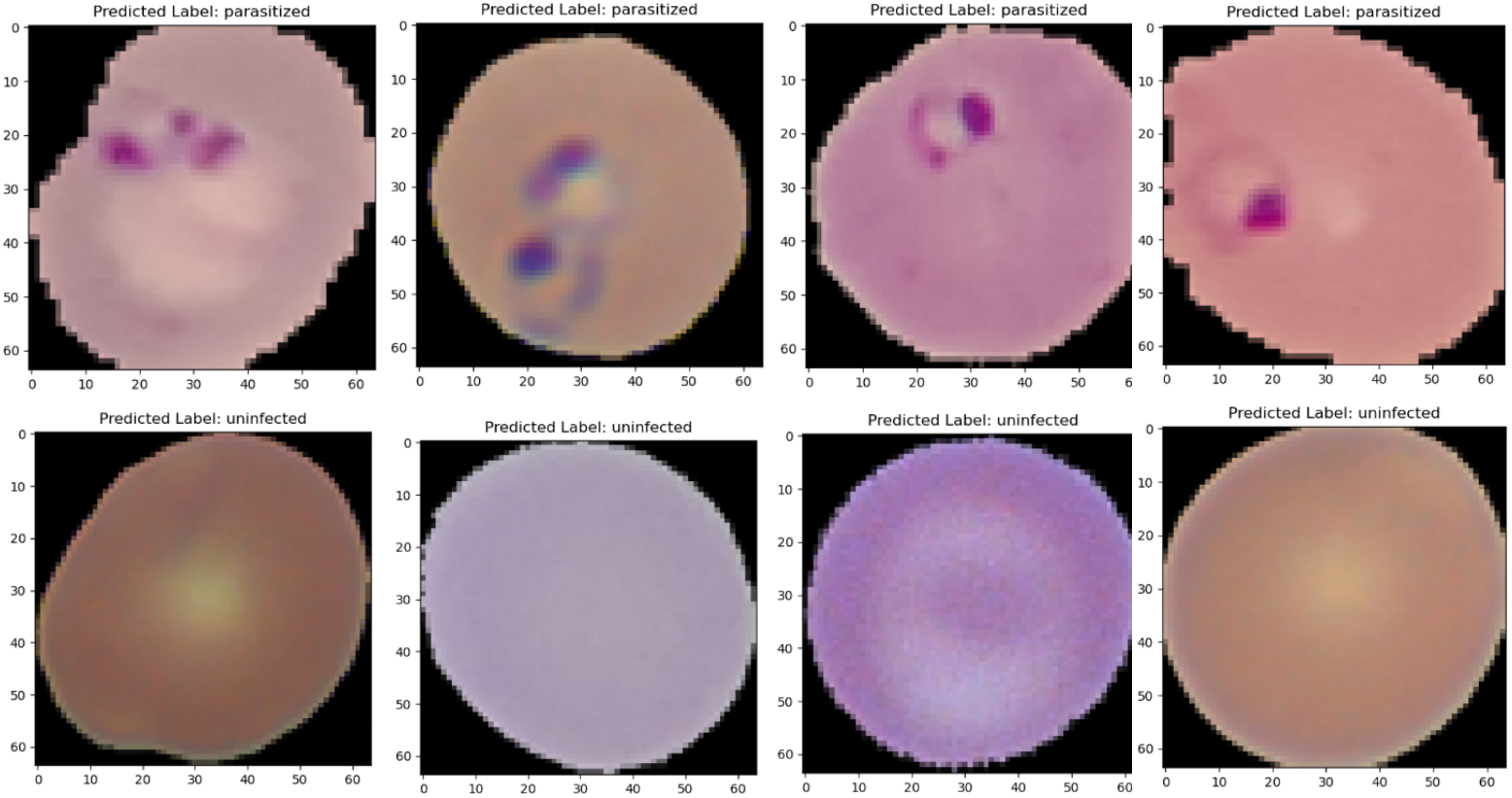
Plasmo3Net model prediction outcome.

The Plasmo3Net comparative study used the four pre-trained models, ResNet50, VGG16, Inception-V3, and AlexNet (Table 2; Figures 4 and 5). Of the three CNN architectures developed, the 13-layer model with three convolutional layers gave the best metrics (Supplementary Figure 1). The two other CNN models were a 16-layer architecture with four convolutional layers and a 19-layer architecture with five convolutional layers. It was observed that the 16-layer performed better than the 19-layer architecture (Table 2). The accuracy, precision, F1 score, and specificity of the 16-layer model were 97.7%, 97.8%, 97.7%, and 97.9%, respectively, compared to the 19-layer architecture metrics: 97.3%, 95.9%, 97.4%, 95.8%, respectively. However, the 19-layer architecture’s recall (98.9%) was higher than that of the 16-layer architecture (97.6%)

**Table 2:** Model performance metrics.

| Model | Accuracy | Precision | Recall | F1 score | Specificity | Sensitivity |
| --- | --- | --- | --- | --- | --- | --- |
| Inception-V3 | 51.3 | 51.3 | 52.4 | 51.9 | 50.2 | 52.4 |
| AlexNet | 91.3 | 93.3 | 89.0 | 91.1 | 93.6 | 89.0 |
| ResNet50 | 97.9 | 97.6 | 98.3 | 97.9 | 97.6 | 98.3 |
| VGG-16 | 97.8 | 98.6 | 96.9 | 97.7 | 98.6 | 96.9 |
| 16-layer CNN | 97.7 | 97.8 | 97.6 | 97.7 | 97.9 | 97.6 |
| 19-layer CNN | 97.3 | 95.9 | 98.9 | 97.4 | 95.8 | 98.9 |
| Plasmo3Net | 99.3 | 99.1 | 99.6 | 99.3 | 99.1 | 99.6 |

The Plasmo3Net had the best performance considering the accuracy, F1 score, recall, and precision (Supplementary Figures 2 and 3). The Plasmo3Net gave the best result among all other models, with a test accuracy of 99.3%. In terms of precision, the VGG-16 architecture performance (98.6%) was slightly higher than the ResNet50 (97.6%). However, the ResNet architecture outperformed the VGG-16 model considering the other three evaluation metrics. The ResNet50 F1 score, recall, and accuracy were 97.9%, 98.3%, and 97.9%, respectively, while the VGG architecture had 97.7%, 96.9%, and 97.8%, respectively.

The Inception V3 gave the lowest performance considering all the evaluation parameters (test accuracy 51.3%, precision 51.3%, recall 52.4%, and F1 score 51.9%). When we compared the four-evaluation metrics, the Plasmo3Net provided superior performance. As a result, the Plasmo3Net outperformed the CNN-based transfer-learning models in recognizing parasitic RBCs, even though it had fewer parameters (Figures 7 and 8). When the performance and the effectiveness of Plasmo3Net were evaluated using the manually curated NIH dataset (Supplementary Figures 4), we observed high-performance metric values. The model obtained an accuracy and precision of 97.6% and 96.9%, respectively. The model had a remarkable F1 score and recall rate of 97.6% and 98.3%, respectively.

**Figure 7.**
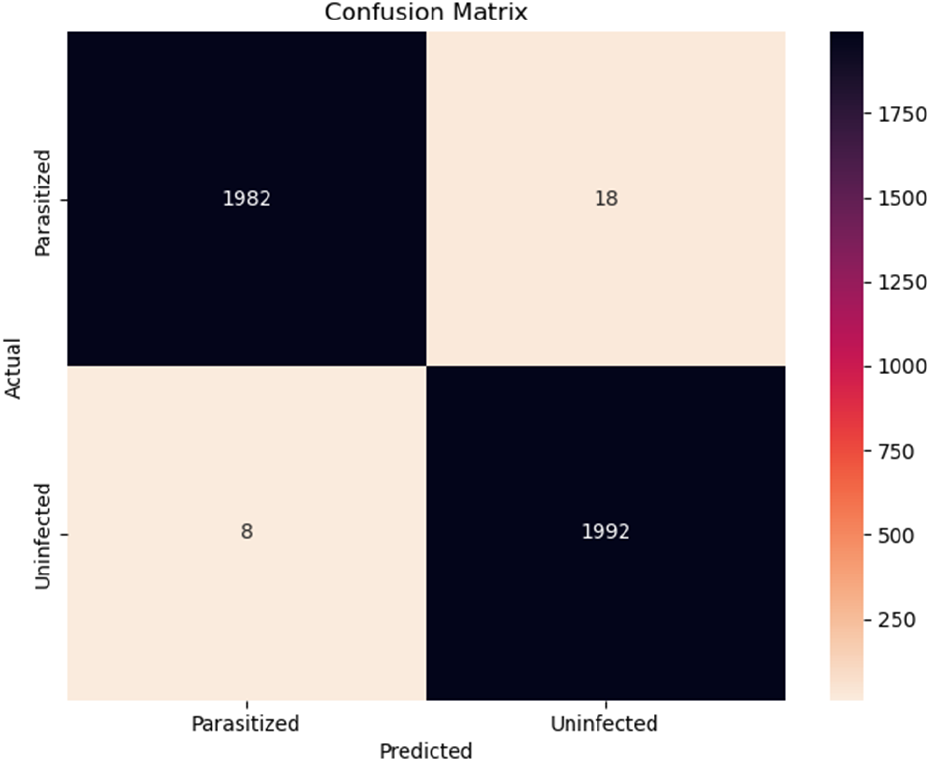
Confusion matrix for Plasmo3Net evaluation

**Figure 8.**
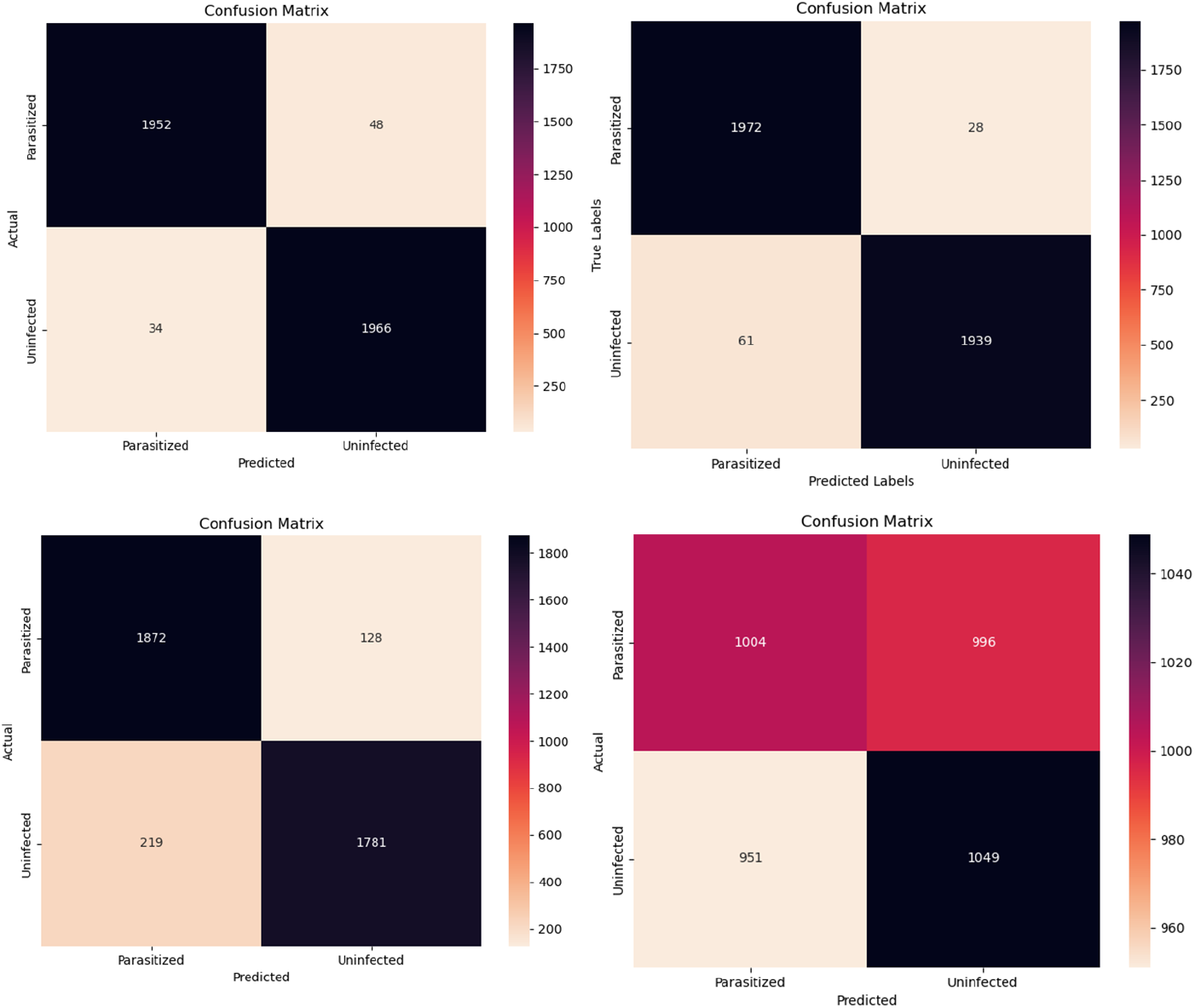
Evaluation of confusion matrices for pre-trained architectures: (A) ResNet50, (B) VGG-16, (C) AlexNet, and (D) Inception V3.

## 4.0 DISCUSSION

Early malaria diagnosis is vital for effective control and eradication. Traditional diagnostic tools often fall short, especially in resource-limited settings where accurate and timely detection is crucial. Integrating artificial intelligence into malaria control strategies offers a promising solution to address these challenges and support elimination efforts. To enhance diagnostic accuracy and speed, this study focused on developing an automated deep learning algorithm based on CNN to detect red blood cells (RBCs) infected with *P. falciparum*.

The CNN algorithm developed in this study, Plasmo3Net with a 13-layer architecture, obtained a very high performance after hyperparameter optimization. Optimization of the batch size led to a significant reduction in the cost function, demonstrating the effectiveness of this hyperparameter tuning. The Plasmo3Net demonstrated superior performance with higher accuracy compared to an 8-layer CNN architecture with the same number of epochs (25) and batch sizes [30]. Furthermore, the 13-layer Plasmo3Net architecture with 3 convolutional layers performed better than the 16-layer architecture with 4 convolutional layers as well as the 19-layer model with 5 convolutional layers. The exceptional performance of the 13-layer Plasmo3Net is likely attributed to its compact architecture [31]. The structure of complex 16- and 19-layer algorithms can lead to models becoming overly tailored to the training set, a phenomenon known as overfitting. In such cases, the models learn not only the underlying patterns in the training data but also the noise and outliers, which compromises their ability to generalize to new, unseen data. This over-specialization could result in a decline in model performance when applied to datasets beyond the training environment, thereby limiting its practical utility [32, 33]. In addition, the additional layers in the 16-layer and 19-layer models may not have effectively captured unique features or contributed meaningfully to the classification process, potentially reducing their overall performance. In the 19-layer model, a very high learning rate could lead to issues with gradient descent not converging, potentially impacting its effectiveness. [34]. However, our simple 13-layer model successfully extracted sufficient features from the dataset for effective generalization.

Insights into Plasmo3Net’s learning progress revealed a significant decrease in training loss. The model’s accuracy was a result of its high performance throughout the learning phase, demonstrated by both training and validation accuracy, as well as training and validation losses. The Plasmo3Net outperformed a previously customized CNN models which also obtained high accuracy in a ten-fold cross-validation [35, 36]. Moreso, the performance metrics of Plasmo3Net were superior to other pre-trained models selected for comparison. ResNet50 [35], VGG16 [36], Inception-V3 [37], and AlexNet [36] were selected as baseline models due to their established performance in malaria parasite identification. The superior performance of Plasmo3Net highlights its potential as a reliable tool for malaria parasite identification. Its adaptability as a reference model for training a more diverse red blood cell (RBC) dataset is crucial in the context of global health, where accurate and timely diagnosis significantly influences treatment outcomes.

Moreover, Plasmo3Net demonstrated not only high accuracy but also a high precision rate. This indicates that the model effectively reduced false positives while accurately identifying positive cases. The model’s precision likely resulted from its sophisticated data preprocessing techniques, as well as its architecture, which enhanced feature extraction and decision-making. The precision of the Plasmo3Net was higher than previously reported models [30, 38, 39]. However, it is essential to consider the implications of high precision in the context of recall [40]. A model optimized for precision may sometimes trade-off recall, potentially leading to missed positive cases [41]. Interestingly, our model achieved a high recall despite this potential trade-off. In addition, our proposed model achieved the highest F1 score, indicating its proficiency in accurately identifying true positive instances while effectively reducing false positives and false negatives. The Plasmo3Net F1 score surpassed previous 16-layer model architectures containing 4 convolutional layers after batch-size optimization to 32 [35, 42].Good performance measures were also obtained by this model when tested on an external dataset.

## Limitations and Future Directions

Our model showed strong performance in detecting *P. falciparum*-infected RBCs, but several limitations should be acknowledged to properly contextualize this finding. For example, the model was not trained on different *Plasmodium* species and did not account for the parasite’s various life-cycle stages. This limitation could impact the model’s accuracy and sensitivity in identifying specific malaria types, as different life-cycle stages exhibit unique morphological characteristics. To address this limitation, future research could focus on developing a convolutional neural network (CNN)-based multiclass classification model capable of performing speciation and identifying different life-cycle stages. Such advancements would be crucial for accurate diagnosis, effective treatment, research, and policy formulation. Nevertheless, this study has demonstrated the potential of CNNs in advancing image-based analysis and diagnosis of malaria.

### CONCLUSIONS

This study has developed Plasmo3Net, a CNN-based deep learning algorithm for classifying red blood cells (RBCs) infected with *P. falciparum*. The model demonstrated superior performance compared to previous models, highlighting the importance of hyperparameter optimization, such as batch size and epoch count, and achieving very high-performance metrics. This has laid the groundwork for future investigation and improvement in malaria diagnosis.

## CONFLICT OF INTERESTS

The authors declare no conflict of interest.

## FUNDING

Kolapo M. Oyebola was supported by the European and Developing Countries Clinical Trials Partnership (EDCTP) Career Development Fellowship (TMA2019CDF-2782). The views expressed in this publication are those of the authors and not necessarily those of the European Union.

## AVAILABILITY OF DATA AND MATERIALS

The dataset supporting the findings of this article are within the manuscript and its Supplementary File.

## ACKNOWLEDGEMENTS

The authors appreciate the study participants and Ijede community leaders for their cooperation during the field collection. Special gratitude to the students and staff of the Center for Genomic Research in Biomedicine (CeGRIB), Mountain Top University for their clerical support during manuscript drafting.

## AUTHORS’ CONTRIBUTIONS

Kolapo M. Oyebola and Afolabi J. Owoloye conceived and designed the study. Kolapo M. Oyebola, Funmilayo C. Ligali, Oluwagbemiga O. Aina and Afolabi J. Owoloye collected field samples. Kolapo M. Oyebola and Afolabi J. Owoloye implemented data analysis, including deep learning algorithms. Kolapo M. Oyebola and Afolabi J. Owoloye drafted the manuscript. All authors read, revised and approved the final manuscript.

## Notes

### Competing Interest Statement

The authors have declared no competing interest.

