## Supplemental File for "Plasmo3Net: A Convolutional Neural Network-Based Algorithm for Detecting Malaria Parasites in Thin Blood Smear Images"

### Slide 1
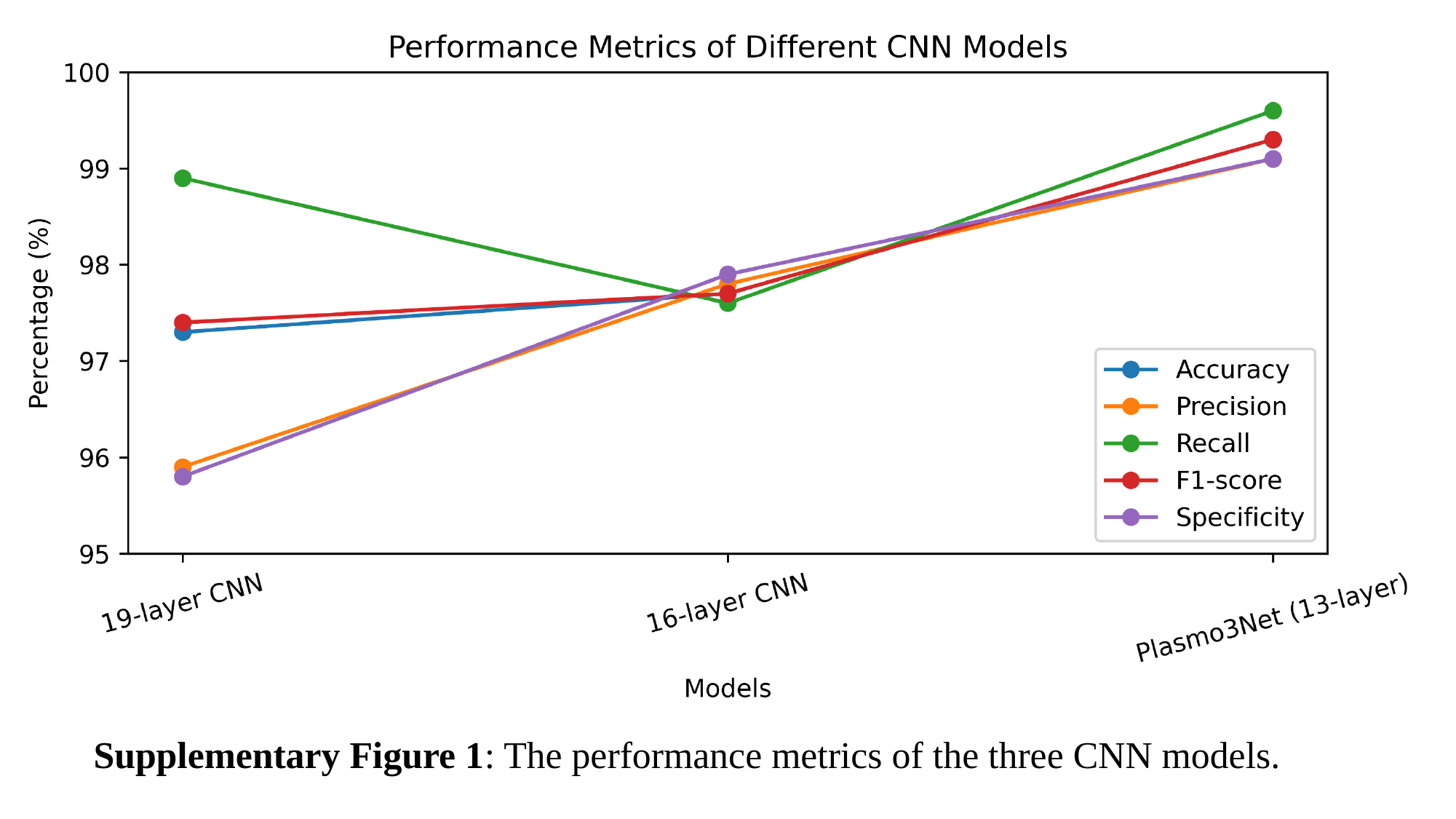

Supplementary Figure 1: The performance metrics of the three CNN models.

### Slide 2
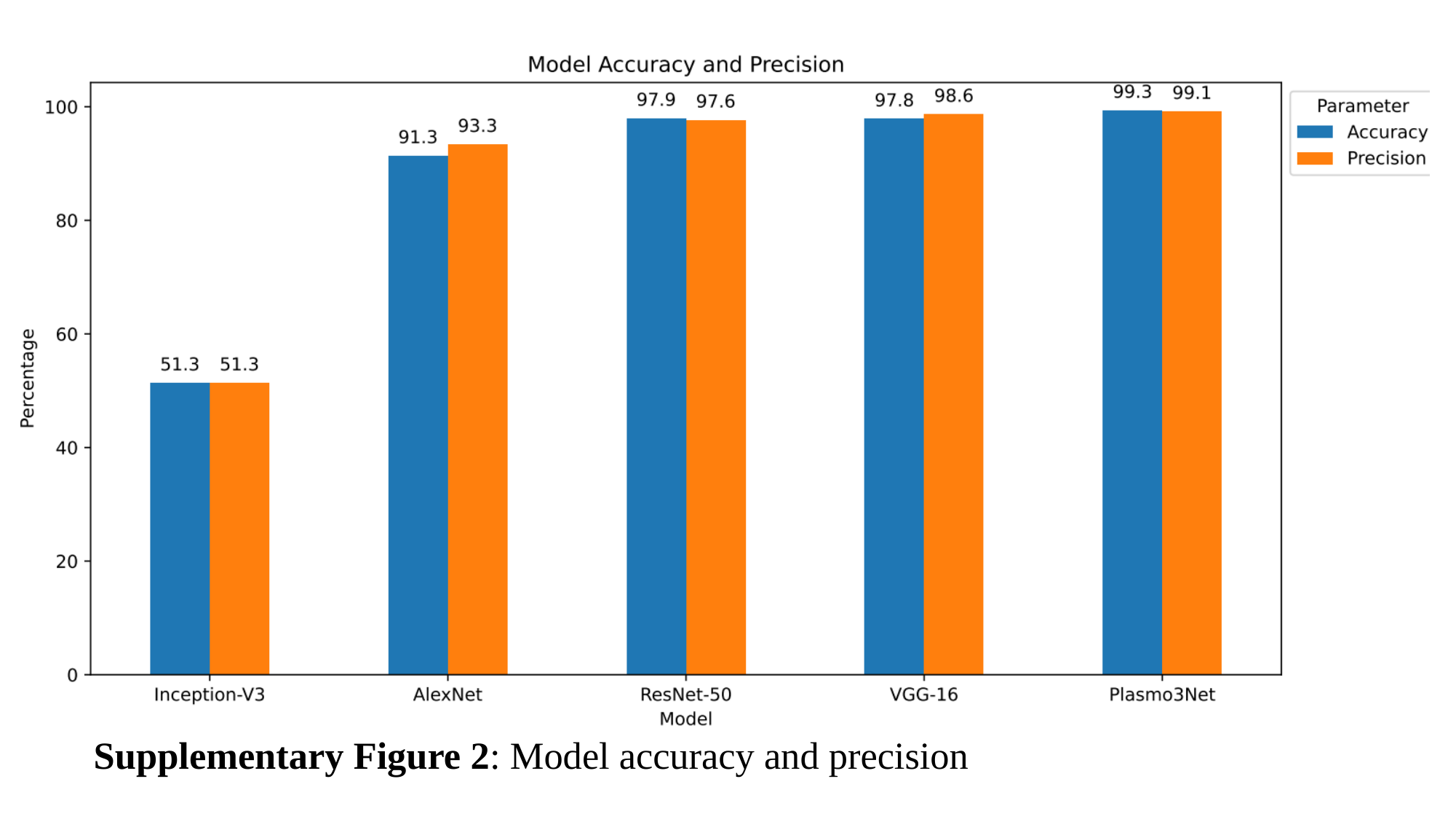

Supplementary Figure 2: Model accuracy and precision

### Slide 3
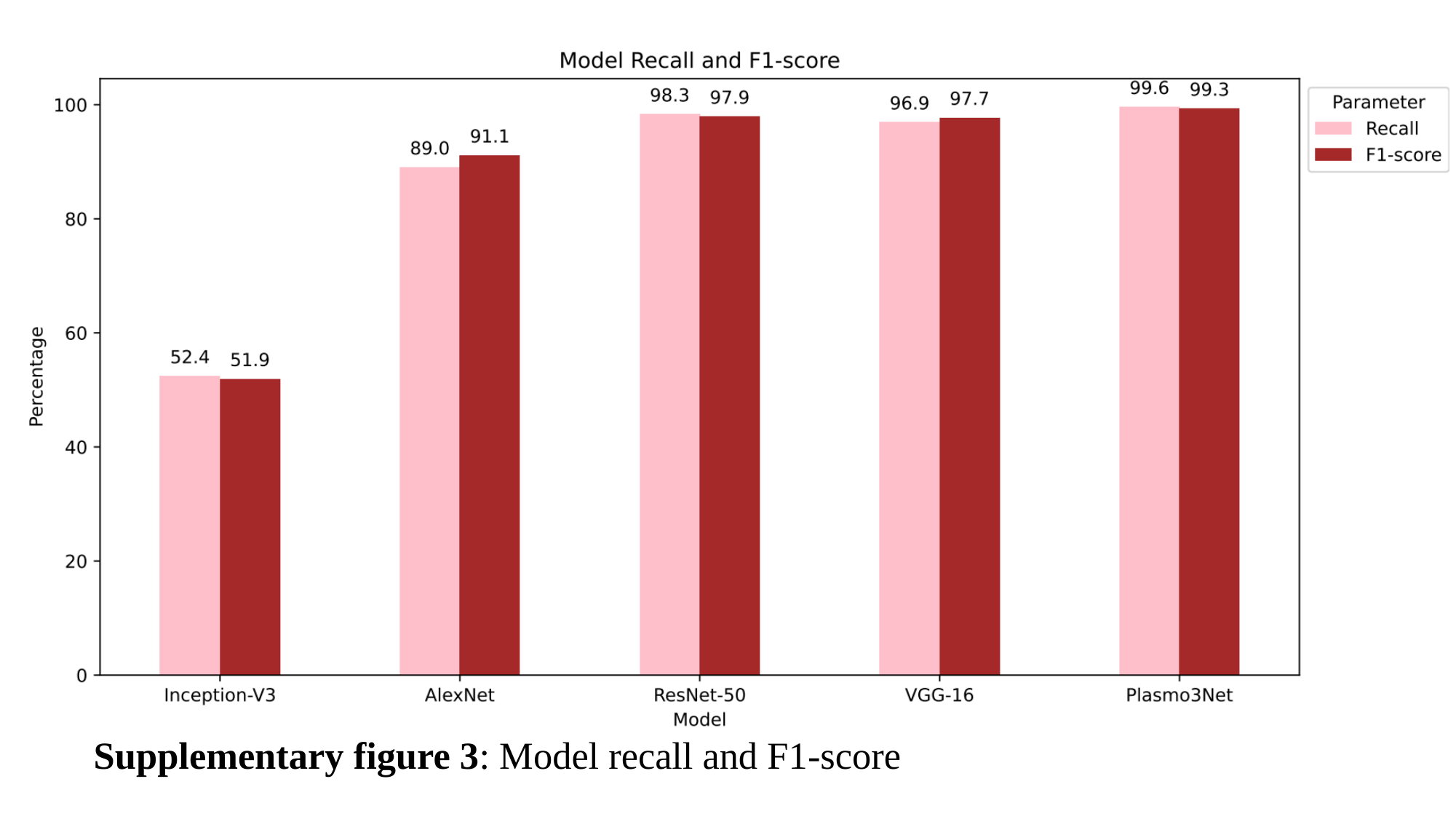

Supplementary figure 3: Model recall and F1-score

### Slide 4
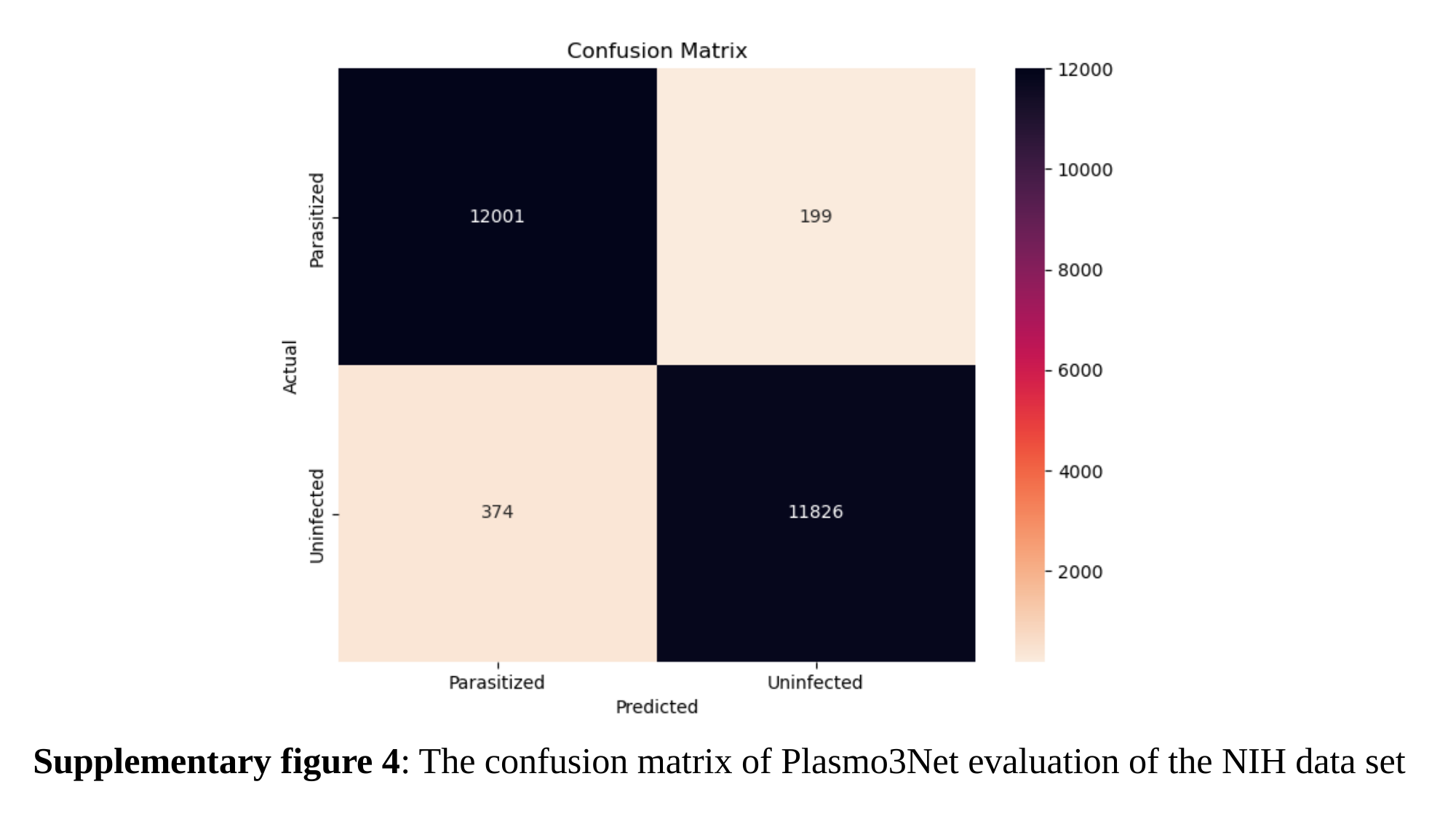

Supplementary figure 4: The confusion matrix of Plasmo3Net evaluation of the NIH data set
